# The Impact of Instrumental Music on Dual-task Interference in a Simulated Driving Environment

**DOI:** 10.1101/2024.12.05.627086

**Authors:** Elahe Khadiv, Ali Salehi Sahlabadi, Reza Hashemirad, Mahshid Namdari, Mojtaba Abbaszadeh

## Abstract

Numerous drivers engage in both listening to music and undertaking secondary tasks while driving. Secondary tasks have been shown to have a negative impact on driving performance. However, the effect of music on driving performance can vary, either improving or impairing it. Nevertheless, the influence of background music on dual-task interference remains uncertain. In this study, we aimed to investigate the impact of instrumental background music on dual-task performance within a simulated driving environment. Twenty-four participants performed a color-discrimination task followed by a driving lane-change task. The results indicated that music enhanced accuracy in the lane-change task without a corresponding increase in reaction time (RT), both in single- and dual-task conditions. Additionally, when the time gap between the two tasks was 600 ms, RT for the lane-change task decreased with high music tempo. However, RT for the lane-change task did not exhibit any changes in response to music tempo when the onset time between the two tasks was short (100 ms), as well as in the single-task condition. Notably, participants’ perceived workload did not significantly differ across the various task conditions. These findings emphasize the potential of background music to improve dual-task performance, particularly when considering the temporal gap between tasks.

## Introduction

In today’s fast-paced world, the prevalence of dual-tasking while driving has become increasingly common. With the widespread use of smartphones and in-vehicle technology, drivers often find themselves juggling multiple activities behind the wheel. However, the effect of engaging in secondary tasks on driving performance has raised significant concerns for road safety. Numerous scientific studies have delved into this subject, shedding light on the detrimental impact of dual-tasking on driving abilities (Abbas-Zadeh, Hossein-Zadeh, & Vaziri-Pashkam, 2021; Hibberd, Jamson, & Carsten, 2013; Levy, Pashler, & Boer, 2006; Strayer, Drews, & Johnston, 2003). These studies consistently demonstrate that attempting to divide limited cognitive resource between driving and other tasks can lead to impaired performance and an increased risk of accidents. Furthermore, it is well-documented that a significant majority of drivers not only engage in performing secondary tasks (Dibben & Williamson, 2007; Insurance, 2000) but also listen to music while driving (Braitman & McCartt, 2010; Huemer, Schumacher, Mennecke, & Vollrath, 2018; Huemer & Vollrath, 2011). However, the impact of background music on driving performance during concurrent secondary task engagement remains an area that requires further investigation.

Studies examining the influence of music on cognitive tasks have yielded intriguing results, revealing a range of effects, both positive and negative, on performance (Harmon, Troester, Pickwick, & Pelosi, 2009; Patston & Tippett, 2011; Schellenberg, 2005). Furthermore, prior research has broadly investigated the ramifications of music on driving performance (Husain, Thompson, & Schellenberg, 2002; Wen, Sze, Zeng, & Hu, 2019). Some studies have shown that listening to preferred music can enhance driver alertness and mood, resulting in improved RT and overall performance (Cassidy & MacDonald, 2010; Ünal, Steg, & Epstude, 2012; Van Der Zwaag et al., 2012). However, Warren Brodsky and colleagues (Brodsky, 2001; Brodsky & Slor, 2013; Young, Mitsopoulos-Rubens, Rudin-Brown, & Lenné, 2012) caution against potential distractions posed by music, as it can divert attention from the road and compromise critical cognitive processes necessary for safe driving. Previous investigations have also compared the effects of instrumental and vocal music on cognitive performance (Crawford & Strapp, 1994; Shih, Huang, & Chiang, 2012; Waters, 2013). Generally, these studies indicate that vocal music exerts a significant negative impact on attention and performance compared to instrumental music. Collectively, the findings of previous research suggest that the influence of music on performance, particularly in the context of driving, varies depending on the type of music. Preferred and instrumental music conditions may enhance driving performance but could also potentially have a negative impact on performance.

The tempo is specific aspect of music that has garnered attention in the recent studies. Different tempos can elicit varying cognitive and emotional responses (Kellaris & Kent, 1993; Oakes & North, 2006; Shalev, Bauer, & Nobre, 2019). Research investigating the impact of music tempo on driving performance has provided valuable insights. Studies consistently suggest that tempo plays a crucial role in influencing driver behavior. For instance, research by Cassidy and MacDonald (2010) found that higher tempo music tends to lead to increased arousal levels and faster RT, potentially enhancing alertness and responsiveness during driving. However, this effect may vary based on individual preferences and driving conditions. Conversely, slower tempo music has been associated with a calming effect, potentially reducing stress levels and promoting a more relaxed driving environment (Li et al., 2010). Nevertheless, caution must be exercised in selecting music with an appropriate tempo, as excessively fast rhythms may lead to overstimulation and decreased focus, while overly slow tempos could induce drowsiness or sluggish responses. Overall, the tempo of music can significantly influence driving performance, highlighting the importance of considering this factor for a safer and more efficient driving experience.

Limited studies have delved into the combined impact of music and secondary tasks while driving (Bellinger, Budde, Machida, Richardson, & Berg, 2009; Millet, Ahn, & Chattah, 2019). Notably, Bellinger et al., (2009) scrutinized the influence of music and cellphone use during driving. Their results indicated that, in general, participants exhibited higher RT when engaged with their cellphone, regardless of whether music was present. This suggests that music did not significantly alter driving RT, whether or not a cellphone was in use. Another study by Millet et al. (2019) compared the effects of background music and cellphone use on driving performance. While this study did not explore the simultaneous effects of cellphone use and background music on driving performance, it demonstrated that cellphone conversations had a more pronounced negative impact on driving performance compared to background music. However, these studies did not systematically control for the onset of driving events and cellphone use. Consequently, they were unable to investigate how music affects the magnitude of dual-task interference while driving.

The Psychological Refractory Period (PRP) paradigm stands as a widely employed method for studying dual-task interference. In this paradigm, participants are presented with two distinct stimuli in rapid succession, each acting as the cue for a unique choice RT task with specific response options. The interval between the onsets of these two stimuli, known as the stimulus onset asynchrony (SOA), is manipulated across trials. The RT and accuracy of both tasks are examined as a function of SOA. Typically, the performance of the first task remains largely unaffected by SOA, while the performance of the second task deteriorates with a decrease in SOA. Previous studies suggest that the performance decline in dual-task conditions arises from the competition between the two tasks for access to limited cognitive resources, as well as the presence of bottlenecks in response selection and execution (Pashler, 1994; Sigman & Dehaene, 2005).

Most prior dual-task studies have focused on simple two-choice tasks, with only a handful of investigations utilizing the PRP paradigm in a driving context (Hibberd et al., 2013; Levy et al., 2006). We have designed a PRP paradigm and replicated the primary findings of previous studies within a simulated driving environment in our prior research (Abbas-Zadeh et al., 2021; Abbaszadeh, Hossein-Zadeh, Seyed-Allaei, & Vaziri-Pashkam, 2023; Hashemirad, Vaziri-Pashkam, & Abbaszadeh, 2023). While our driving PRP paradigm is not an exact match to real-world driving, we think it captures the core features of a lane-change scenario with systematic control over the timing of task onsets, akin to a classic PRP paradigm. It emulates the need for sequential motor movements in driving tasks, mirroring situations where drivers must execute a series of precise actions. For instance, making a turn requires rotating the wheel in one direction and then adjusting it in the opposite direction to straighten the vehicle. This distinctive feature increases the complexity of the driving task compared to simpler artificial tasks. Time sensitivity is a crucial factor within the simulator, mirroring real-world driving situations where prompt reactions are essential. Unlike simple two-choice tasks, most driving actions come with a built-in time limitation. This temporal constraint significantly influences participant behavior compared to tasks without such restrictions. In this dual-task paradigm, the driving task takes precedence over secondary tasks, acknowledging the intrinsic importance of each. Unlike simple two-choice tasks where tasks may carry equal weight, our simulator recognizes the primary focus on driving, influencing participant behavior accordingly. The simulator recreates a continuous, immersive driving environment, complete with distracting elements such as the road, roadside features, and participant-vehicle interactions. Additionally, it incorporates car components like the dashboard and odometer, which can either enhance responses or act as potential distractions, ultimately shaping participant behavior.

In this study, our objective was to examine the impact of background music with varying tempos on dual-task performance within a PRP paradigm implemented in a simulated driving environment. Specifically, participants concurrently undertook a lane-change task alongside a color discrimination task, either in a dual-task configuration or separately as single-task conditions. This was conducted under conditions involving both the absence and presence of background music, featuring compositions with low, medium, and high tempos. The background music consisted of instrumental tracks selected based on participants’ preferences. The results unveiled that executing an immediate lane-change after the color stimulus (short SOA) led to the increase of RT when music was present. However, when there was a longer interval between the color discrimination and lane-change tasks (long SOA), both RT and accuracy demonstrated improvement. Notably, music improved and mitigated the accuracy costs in the dual-task condition for color discrimination.

## Materials and Methods

### Participants

The study comprised a total of 24 participants, with an equal distribution of males and females, aged between 20 and 30 years (mean age: 21.6 ±1.52 years). None of the participants reported being regular gamers, engaging in computerized games for more than two hours per month over the past two years, and they had no known history of neurological diseases. All participants had a driver’s license, normal or corrected-to-normal vision, and no color blindness. Informed consent was obtained from all participants, signifying their voluntary agreement to take part in the study, and they were appropriately compensated for their invaluable contributions. The current research was reviewed and approved by the ethics committee of Shahid Beheshti University of Medical Sciences (ethics code: IR.SBMU.REC.1401.094) in accordance with the Declaration of Helsinki.

### Dual-task paradigm

The dual-task paradigm incorporated both a driving lane-change task and a tone discrimination task. The driving environment was designed using the Unity 3D game engine. The driving scenario encompassed a highway featuring an endless expanse of lanes on both sides, devoid of any left or right turns, as well as inclines or declines. The lanes maintained a straight trajectory, devoid of curves. The car’s movement remained consistently parallel to these lanes, and utilizing the button controls prompted the car to shift between lanes to the right or left. The driving stimulus comprised two rows of traffic cones, each row consisting of three cones (Figure 1). During each trial, the cones unexpectedly appeared on both sides of one of the lanes, requiring participants to immediately steer the vehicle towards the lane and navigate through the cones. The spacing between the two rows of cones allowed the vehicle to traverse through them effortlessly. The cones consistently appeared on the lane directly to the left or right of the driving lane, prompting participants to make a single lane change per trial. Participants were instructed to press and hold the appropriate key to guide the vehicle between the two rows of cones and subsequently release the key once the vehicle was appropriately positioned.

**Figure 1.**
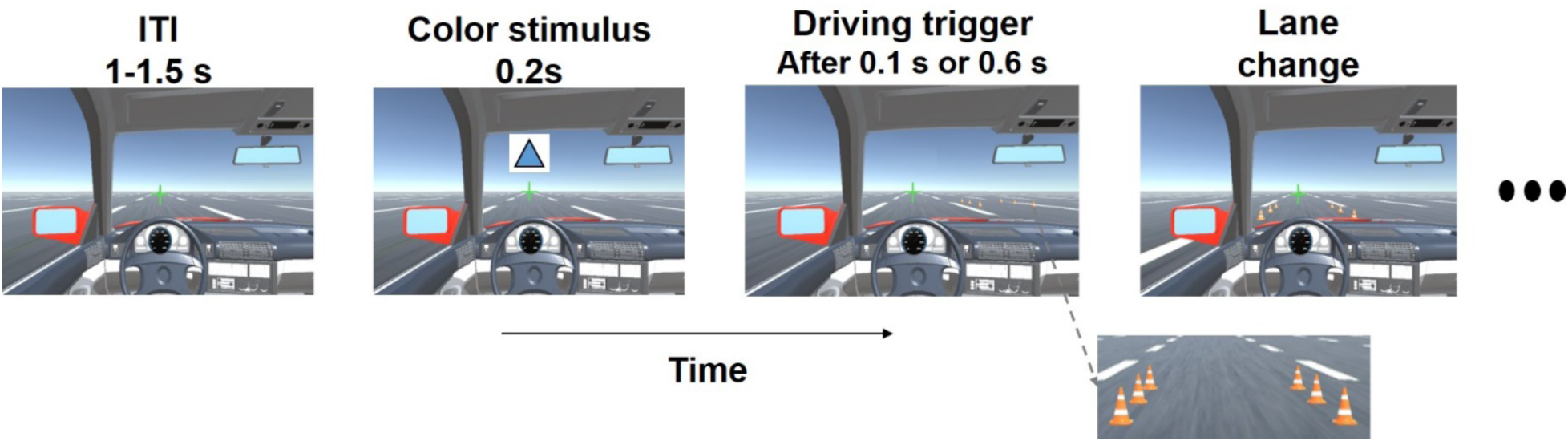
Illustration of a sample trial in the dual-task paradigm. The intertrial interval (ITI) ranged from 1 to 1.5 s, followed by a 200 ms color stimulus. The driving trigger was presented either 100 or 600 ms after the color stimulus. Participants were required to discriminate the color of the geometric stimulus (blue vs. red) immediately after its presentation, followed by the lane-change task upon the presentation of the driving stimulus (represented by two rows of cones).

The vehicle maintained a consistent speed of 80 km/h. The cones were consistently positioned at 44.4 m from the car. Given the speed of 80 km/h, the car would cover this distance in 2 seconds. The width of each lane measured 3 m and executing a lane change at this constant speed required 1.1 sec. To prevent collision with the cones, participants had a window of 0.9 sec to initiate a lane change once they spotted the cones. Participants were directed to change lanes in every trial, even in cases where a delay made collision inevitable. Otherwise, in the subsequent trial, the cones would be positioned two lanes away from them. All participants adhered to this instruction and carried out lane changes in all trials.

The lane change speed was adjusted to ensure a gradual and smooth transition, avoiding the need for constant maneuver corrections by participants. Participants were advised to remain in the middle of their lane unless a lane change was required. While participants had the option to initiate lane changes through a sequence of individual button presses, they were guided to execute a lane change using a single button press-hold-release action, a skill they acquired during training sessions.

The RT for driving was defined as the time from the onset of the driving trigger to the first button press that initiated the lane change. A lane change was deemed successful when the participant pressed and held the key corresponding to the cone’s location until their vehicle was positioned between two cones, at which point they released the key. If the participant mistakenly pressed the right key instead of the left key, even momentarily, the trial was categorized as an error due to an initial incorrect decision. The RT for these error trials was calculated from the trigger onset to the initial incorrect key press. Notably, we did not encounter any instances in which subjects failed to correct their course. To provide real-time feedback in the event of a collision with the traffic cones, the fixation cross underwent 100 ms of jitter. The jitter induced a displacement of 0.1 degree of visual angle, and the direction of this displacement changed every 25 ms.

The experiment commenced with an initial speed set at a constant rate of 80 km/h. Throughout the trial, participants adjusted the car position in either the right or left lanes by using their left hand’s middle and index fingers to press keys, respectively.

For the color discrimination task, a single image with a white background and various geometric shapes in two colors (red and blue) was presented for 200 milliseconds at the center of 2-degree elevation above the crosshair (Figure 1). The size of color stimuli was 2.5 degrees of visual angle. Participants used the “x” and “z” keys on the computer keyboard to perform the color discrimination task of the geometric shapes. The color stimulus for each trial was randomly chosen from a pool of 10 images featuring two distinct colors (red or blue) and various geometric shapes. If participants gave an incorrect response, the cross mark changed from steady green to red. Each trial’s start time corresponded to the color stimulus presentation, while its end was linked to the car’s rear end reaching the set of traffic cones.

The study included both dual-task and single-task conditions. In the dual-task trials, two tasks were presented with either a short (100 ms) or a long (600 ms) SOA. In the single-task trials, either the lane-change task or the color discrimination task was presented alone. There was a total of eight dual-task conditions, combining two SOAs, two lane-change directions for driving (shift right/shift left), and two colors (blue/red). Additionally, there were four single-task conditions, encompassing two lane-change directions and two colors. To amplify the impact of dual-task interference on the lane-change task, the order of task presentation in the dual-task conditions was fixed: the color discrimination task was consistently presented first, followed by the lane-change task. This design was derived from findings in our prior behavioral study and aligned with earlier dual-task paradigms, which demonstrated that the second task is more susceptible to interference.

Each participant completed four blocks of the experiment, including three blocks with background music playing at three different tempos (low, medium and high) and one block without music. All blocks were identical and consisted of three separate runs. The order of blocks was counterbalanced across participants. Each run of every block included 48 trials, with 16 single-task trial (8 single lane-change and 8 single color discrimination) and 32 dual-task trials (16 short SOA and 16 long SOA), displayed in a completely random order to prevent any bias among individuals. Each driving run lasted approximately 210 seconds.

Before conducting the primary experiment, every participant undertook two training sessions in the driving simulator, which mirrored the structure of the main experiment runs. These training sessions aimed to ensure the participants’ proficiency with the simulator controls and their comprehension of how their actions influenced the simulated vehicle’s path. To progress to the primary experimental runs, participants needed to meet a performance threshold of 80% or higher in these training sessions. This requirement was implemented to guarantee that all participants shared a comparable level of previous driving simulator exposure and were capable of accurately executing the task during the main experiment.

### Background music

To determine the tempo of the selected music, we utilized Virtual DJ 8 software (https://www.virtualdj.com) and categorized the songs into three different categories. Participants were then asked to vote online for a variety of pre-prepared music pieces. The piece that received the highest number of votes was chosen as the selected music for each category, which was considered the preferred music for the participants. In the low tempo category, Gavina (Artist) with a tempo of 64.1 received the highest number of votes. In the medium tempo category, Quiver (Artist) with a tempo of 113.2 was selected. Finally, for the high tempo category, Mozart Beyoglunda was chosen. During the experiment, music was played through a speaker positioned behind the monitor, at one meter from the participants. Subsequently, the sound level was measured using a RION NA-26 sound level meter at the participants’ location. To ensure accurate listening, the device was set to the ‘slow’ mode, and the played music had a sound level ranging from 40 to 60 decibels at various tempos and over time.

### Mental workload

Driver Assessment Load Index (DALI) checklist (Pauzié, 2008) was used to investigate the mental workload of participants. DALI is subjective evaluation of mental workload that participant percepts during the perform experiment. DALI consist of seven dimensions: 1) global attention, 2) visual demand, 3) auditory demand, 4) tactile demand, 5) time pressure, 6) stress, and 7) interference. Participants would give a score between 1 (very low) to 5 (very high) to each question. To assess and compare the perceived mental workload among different study blocks, participants were asked to complete the DALI checklist at the end of each block.

### Data analysis and statistical tests

To examine the differences between the music and non-music conditions, paired sample t-tests were conducted. One-way repeated measures ANOVA was employed to assess the effects of tempo on both RT and accuracy in both tasks. Furthermore, two-way repeated measures ANOVA was utilized to explore the interactions between music (music and non-music) and task conditions (long SOA, short SOA, and single task), as well as between music tempo (low, medium, and high) and task conditions. In addition, the mental workload across experiment blocks for each dimension of DALI was compared using repeated measures ANOVA.

## Results

### Comparison of Single- and Dual-Task Performance

Before investigating the music effect on dual-task performance, the RT and accuracy of both tasks were compared across task conditions in non-music block. In the dual-task conditions, where the lane-change task was presented second, the RT and accuracy for this task was significantly influenced by the timing and sequencing of tasks (Figure 2A & 2C; RT: F_2,46_ = 109.74, p < 0.0001, η^2^ = 0.82; Accuracy: F_2,46_ = 11.95, p < 0.001, η^2^ = 0.34). In particular, in the short Stimulus Onset Asynchrony (SOA) condition, where the tasks were presented in close temporal proximity (100 ms), the RT and the accuracy for the lane-change task were significantly higher and lower, respectively, compared to both the long SOA (RT: t_23_ = 14.501, p < 0.0001, η^2^ = 2.03; Accuracy: t_23_ = 5.65, p < 0.001, η^2^ = 0.85) and single lane-change (RT: t_23_ = 8.57, p < 0.0001, η^2^ = 1.68; Accuracy: t_23_ = 2.66, p = 0.014, η^2^ = 0.66) conditions. This indicates that the short SOA led to slower responses in the lane-change task compared to the other conditions.

**Figure 2.**
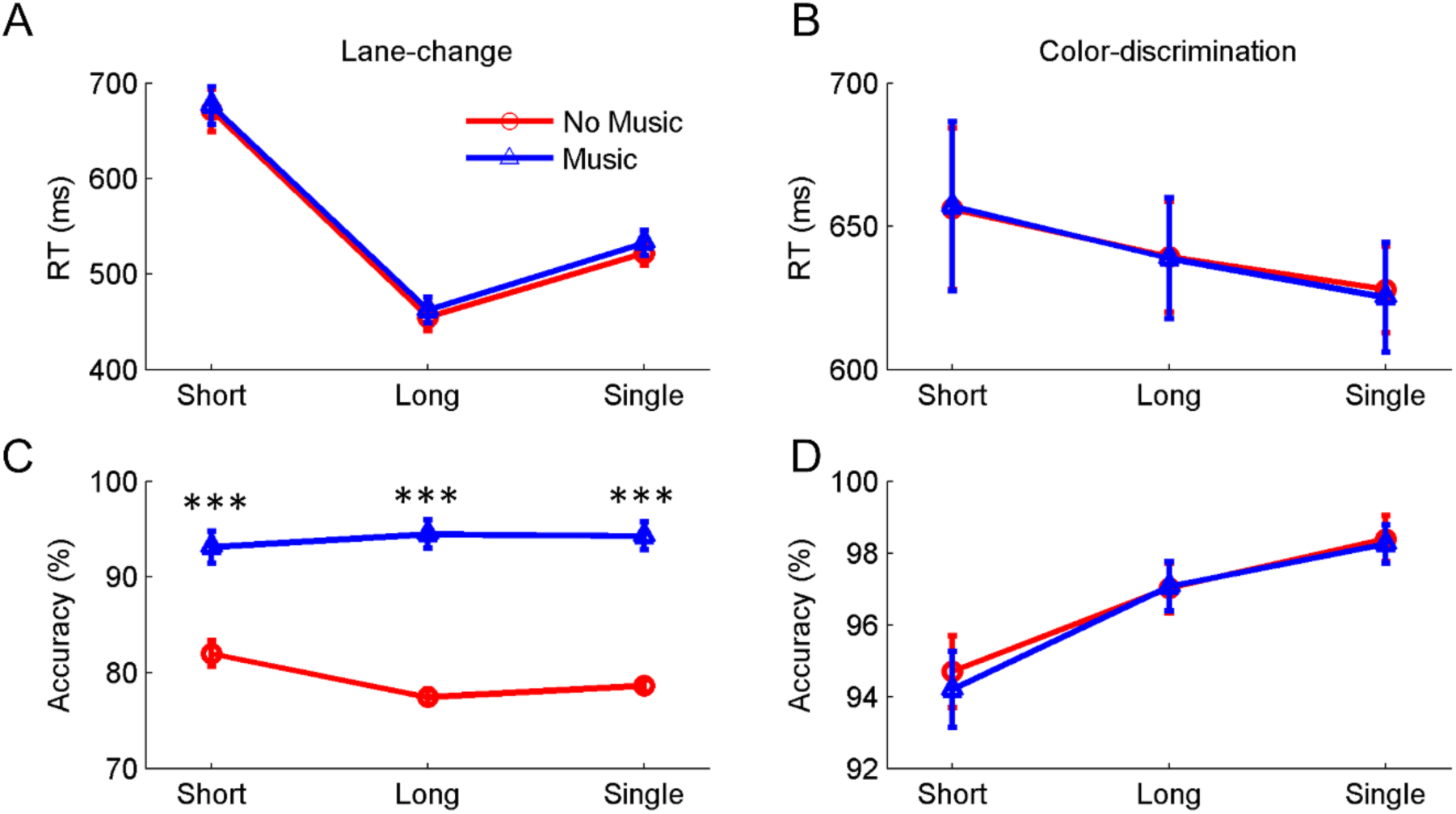
Effect of music on lane-change task. A & B) The RT and accuracy of the lane-change task for short SOA, long SOA and single-task, respectively, in the music (blue) and non-music (red) conditions. C & D) The same format as A & B but for the color discrimination task. In all panels, errorbars show standard error of mean and stars show a significant difference between task conditions for each condition (*** < 0.001).

In the other hand, for the color discrimination, which was always presented before the lane-change task in dual-task conditions, there were no significant differences in RT for this task across single-task, short SOA, and long SOA conditions (Figure 2B; F_2,46_ = 1.54, p = 0.32, η^2^ = 0.04). However, a significant difference in accuracy of the color discrimination task was observed between the task conditions (Figure 2D; F_2,46_ = 11.13, p < 0.001, η^2^ = 0.32). Further pairwise analyses showed that the short SOA accuracy was significantly lower than both the long SOA (t_23_ = −2.72, p = 0.012, η^2^ = 0.50) and single-task (t_23_ = −5.62, p < 0.0001, η^2^ = 0.89) conditions. These results suggest that the timing and sequencing of tasks primarily influenced the RT of the second presented task (lane-change), while the first presented task (color discrimination) remained relatively unaffected. However, it is important to note that the accuracy of the color discrimination task was significantly affected in the dual-task conditions. These findings were consistent with our previous studies (Abbas-Zadeh et al., 2021; Abbaszadeh et al., 2023; Hashemirad et al., 2023)

### Effect of Music on Single- and Dual-Task Performance

To investigate the general effect of music on participants’ performance, the mean RT and accuracy were averaged across three tempo conditions (low, medium, and high) and compared with the performance in the non-music condition. The analysis was conducted separately for single-task and dual-task conditions (Figure 2).

The two-way repeated measures ANOVA was conducted to examine the effects of music (music and non-music) and task condition (short SOA, long SOA and single-task) for the lane-change and color discrimination tasks. The results showed that the main effect of music was only significant for the lane-change accuracy (Figure 2C, F1,23 = 385, p < 0.0001, η^2^ = 0.94) and there was no significant effect of music for the RT of lane-change and the RT and accuracy of the color discriminations task. Nonetheless, the main effect of task condition was significant for the RT and accuracy of the lane-change task and the accuracy of the color discrimination task (Figure 2A, 2C & 2D; Fs_1,23_ > 4.86, ps < 0.012, η^2^ > 0.17), but the task condition had no significant on the RT of color discrimination (p = 0.15). In addition, the analyses revealed that the interaction between music and task condition was also significant only for the accuracy of the lane-change task (F_2,46_ = 13.39, p < 0.0001) and was not significant for the RT of both tasks and the accuracy of color discrimination task (ps > 0.77).

To further investigate the music effect on the lane-change accuracy in each task conditions (short and long SOAs and single task), pairwise comparison was preformed between music and non-music blocks separately for each task condition. The results revealed that the background music improved the accuracy of the lane-change task in both dual-task (short and long SOAs) and single-task conditions (ts_23_ > −8.64, ps < 0.0001, η^2^ > 1.64). Taken together, these findings suggests that the background music improves the lane-change accuracy both dual-task and single task conditions without having negative impact on the RT of lane-change and color discrimination.

### Effect of Music Tempo on Single- and Dual-Task Performance

To explore the impact of music tempo on task performance, we conducted a one-way repeated measures ANOVA to examine the effects of tempo (low, medium, and high) on the RT and accuracy of both the lane-change and color discrimination tasks. The analysis was conducted separately for single-task and dual-task conditions across tempo (Figure3).

For the lane-change task, the analyses revealed a significant effect of music tempo on the RT of the lane-change task specifically in the long SOA condition (Figure 3A). The analysis showed that as the music tempo increased, the lane-change RT significantly decreased in the long SOA (F_2,46_ = 3.58, p = 0.028, η^2^ = 0.14). This suggests that higher tempo music facilitated faster responses during the long SOA condition without having any negative impact on the lane-change accuracy. However, it is important to note that the effect of tempo on RT and accuracy was not significant for the short SOA and single conditions of the lane-change task (ps > 0.15, Figure 3A & 3C), except a marginal effect of the accuracy of lane change in the single task condition (F_2,46_ = 2.74, p = 0.075, η^2^ = 0.10). Overall, these findings indicate that while music tempo had a significant impact on RT during the long SOA condition of the lane-change task.

**Figure 3.**
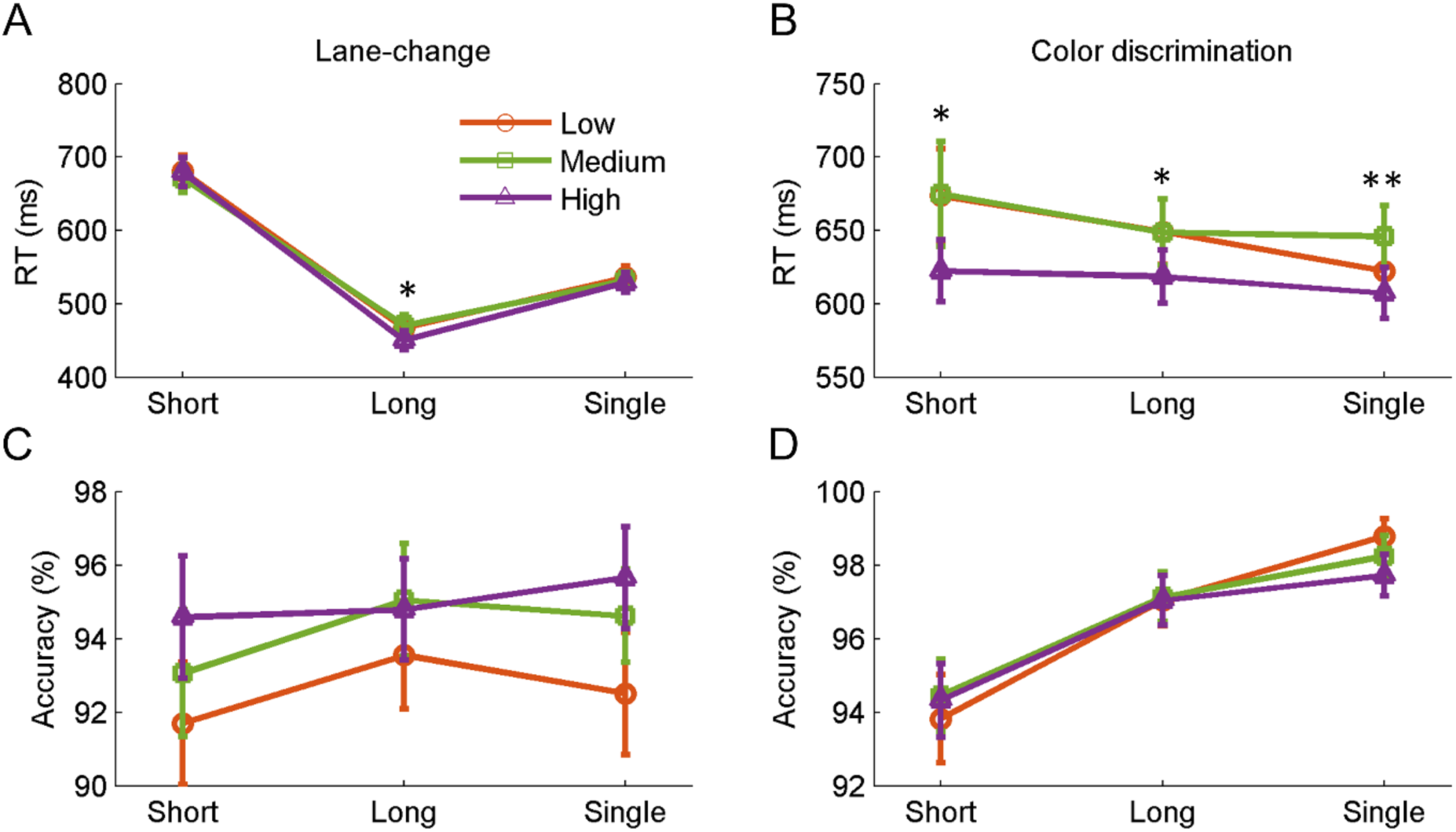
Effect of tempo on lane-change task. A & B) The RT and accuracy of the lane-change task for short SOA, long SOA and single-task across different music tempos (low, medium and high). Orange line is for low temp, green for medium tempo and purple for high tempo. C & D) The same format as A & B but for the color discrimination task. In all panels, errorbars show standard error of mean and stars show a significant difference between task conditions for each condition (* < 0.05 and ** < 0.01).

In contrast, for the color discrimination task, the effect of tempo was found to be significant on the RT for all task conditions (single task, short SOA, and long SOA; Fs_2,46_ > 4.36, ps < 0.018, η^2^s > 0.16; Figure 3B). The results revealed that the RT decreased as the tempo of the music increased in these task conditions. This indicates that participants took shorter to respond to the color discrimination task when exposed to higher tempo music. However, the effect of tempo on the accuracy of the color discrimination task was not significant across all task conditions (ps > 0.40). Therefore, these findings demonstrate that the higher tempo of preferred instrumental music background improved the color discrimination RTs without any change on the accuracy of participants either dual- and single-task conditions.

Additionally, we conducted a two-way repeated measures ANOVA to examine the interaction between task condition (single task, short SOA, and long SOA) and tempo (low, medium, and high) on the performance of both the lane-change and color discrimination tasks. The results revealed that there was no significant interaction effect between task condition and tempo for both the lane-change task (p > 0.11, Figure 3A & 3C) and the color discrimination task (p > 0.20, Figure 3B & 3D). These findings indicate that the combined influence of task condition and tempo did not result in a significant differential effect on the RT or accuracy of either task. This suggests that the individual effects of task condition and tempo on task performance, as previously described, were not further influenced by their interaction.

### Comparison of perceived mental workload in no-music and music conditions

In addition, we assessed the mental workload experienced by participants at the end of each block using the Driver Assessment Load Index (DALI). The results, depicted in Figure 4, showed that the overall mental workload did not significantly vary across the four blocks of the study (non-music, music with low tempo, music with medium tempo, and music with high tempo) for most of the DALI dimensions (Fs_3,69_ < 1.60, ps > 0.19, η^2^s < 0.06). However, a significant difference was observed in the auditory load dimension, where the non-music block had significantly lower scores compared to the music blocks (Fs_3,69_ = 35.04, p < 0.0001, η^2^s = 0.60). This finding is not surprising, as the absence of background music in the non-music block would naturally result in a lower auditory load. Therefore, the observed difference in auditory load between the non-music and music conditions is considered a trivial effect. This additional analysis provides insight into the participants’ perceived mental workload and confirms that the presence of music did not significantly affect the overall mental workload experienced during the task.

**Figure 4.**
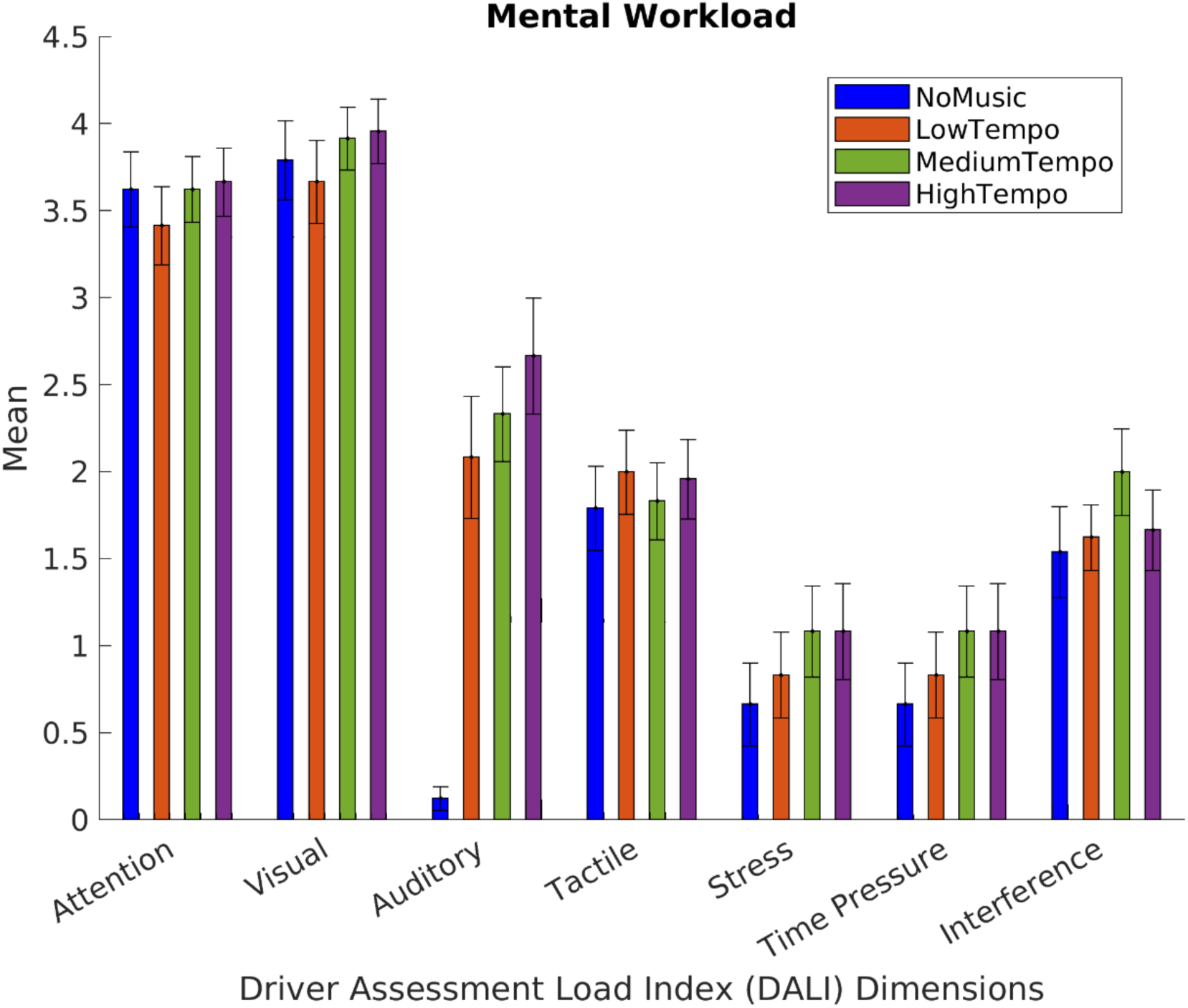
Comparison of Mental Workload Across Experimental Blocks. The x-axis represents different dimensions of the Driver Assessment Load Index (DALI) for four blocks: non-music (black), music with low tempo (dark gray), music with medium tempo (light gray), and music with high tempo (white). The y-axis represents the mean of each dimension across participants.

## Discussion

Our study aimed to investigate the overall impact of preferred instrumental background music on both single- and dual-task performances in a simulated driving environment. The results indicated an enhancement in lane-change performance when music was present compared to non-music blocks, regardless of task complexity (dual-task with short and long SOAs, and single task conditions). Moreover, we observed that an increase in music tempo led to a reduction in lane-change RT specifically in the long SOA, as well as in color discrimination RT across all task conditions (long and short SOAs, and single task). The varying influence of music on the two tasks suggests an order-dependent relationship between music and performance under dual-task conditions, as suggested by LaCroix, Diaz, and Rogalsky (2015). Specifically, the accuracy of lane-change consistently demonstrated improvement when accompanied by background music, regardless of the presentation time of the color discrimination stimulus. However, the RT for lane-change only decreased with a high music tempo when presented a longer interval after the color stimulus. Conversely, the effect of music on the accuracy of the color discrimination task, which was presented second, was not statistically significant, but its RT decreased with an increase in music tempo across all task conditions.

The present study’s results, in line with our previous research, demonstrate that presenting a stimulus prior to the driving task leads to an increase in RT (Abbas-Zadeh et al., 2021; Abbaszadeh, Hossein-Zadeh, Seyed-Allaei, & Vaziri-Pashkam, 2020). This aligns with the findings of Levy et al. (2006) and Hibberd, Jamson, and Carsten (2013), who reported interference effects and task-specific variations in performance during dual-task conditions in simulated driving. Specifically, in the short Stimulus Onset Asynchrony (SOA) condition, where tasks were presented in close temporal proximity, the lane-change task exhibited slower RT compared to both the long SOA and single lane-change conditions. This suggests that dividing attention between two tasks in rapid succession can hinder performance in both tasks. Conversely, in the long SOA condition, where tasks had less temporal overlap, the RT for lane changes was even faster than in the single lane-change condition. This implies that the second task not only failed to increase the RT for lane changes, but also improved it. This improvement may be attributed to the priming effect of the color stimulus on the lane-change task, which was absent in the single lane-change condition where the color stimulus was not presented prior to the task. This finding is in concordance with our recent study (Hashemirad et al., 2023), wherein presenting a tone stimulus 600 ms before the lane-change stimulus resulted in a decrease in RT for lane changes compared to the single lane-change condition.

The comparison of two tasks performance in conditions without and with background music revealed that the background music improves the accuracy of the lane-change task without change on the accuracy color discrimination and RT of both tasks (Figure 2). These findings are consistent with the broader body of literature that suggests music can exert a multifaceted influence on cognitive and motor skills during driving (Cassidy & MacDonald, 2010; Millet et al., 2019). The enhancement in driving accuracy observed in our study can be attributed to several potential mechanisms. Firstly, music, especially when it aligns with the driver’s preferences, has been shown to induce positive emotions and reduce stress and anxiety (Cassidy & MacDonald, 2010). Such emotional regulation may promote a more relaxed yet focused state, which is conducive to maintaining precision in driving tasks. Secondly, music can serve as an auditory background that masks disruptive external noises and distractions, creating a controlled acoustic environment within the vehicle cabin. This can enhance the driver’s ability to concentrate on the road (Ünal, de Waard, Epstude, & Steg, 2013). Additionally, the rhythmic qualities of music may synchronize with the driver’s movements, potentially facilitating coordination and alertness (Ünal et al., 2012). While the exact neurocognitive mechanisms underlying these effects warrant further investigation, our results support the notion that carefully chosen music can serve as a valuable tool to enhance driving task performance, potentially reducing the risk of errors and accidents on the road.

Our study examined the impact of music tempo on task performance across different task conditions. For the lane-change task, we found that music tempo did not have a significant effect on RT or accuracy in all task conditions, except RT in the long SOA. These results align with previous studies by (Navarro, Osiurak, & Reynaud, 2018) and (Millet et al., 2019), which demonstrated music tempo did not has the effect on processing speed and response times in driving. However, it is important to note that these previous studies focused solely on the effect of tempo in single driving conditions, while our study explored the effect in both single and dual-task conditions. Although we observed a decreasing trend in RT with increasing tempo in the long SOA condition. Similarly, there was no significant effect of tempo on the accuracy of the lane-change task. These findings suggest that music tempo did not have a substantial impact on the performance of the lane-change task during dual-task interference conditions.

In our study, we observed a notable difference in the impact of music tempo on the color discrimination task compared to the lane-change task. Specifically, the color discrimination task showed a significant effect of music tempo on RT across all task conditions. This finding supports and extends previous research conducted by Schellenberg, Nakata, Hunter, and Tamoto (2007) and Angel, Polzella, and Elvers (2010), who also reported a similar relationship between music tempo and RT in cognitive tasks, although these studies primarily investigated the effect of tempo on single-task conditions. Our results consistently demonstrated that faster tempo music was associated with faster RT in the color discrimination task. This suggests that the high tempo music may have facilitated participants’ cognitive processing, resulting in quicker and more efficient decision-making. Importantly, the effect of music tempo on task accuracy in the color discrimination task was not found to be statistically significant. This implies that while music tempo influenced the speed of participants’ responses, it did not significantly impact their overall accuracy in discriminating colors.

The assessment of mental workload using the Driver Assessment Load Index (DALI) provided valuable insights into participants’ subjective experience during the study. Our results indicated that the overall mental workload did not differ significantly across the four blocks of the study, including the non-music condition and the music conditions with different tempos. This suggests that the introduction of music during the tasks did not have a substantial impact on participants’ perceived mental workload.

In summary, our study focused on the impact of preferred instrumental background music on simulated driving tasks. We found that having music playing led to improved lane-change performance, regardless of task complexity. Additionally, higher music tempo correlated with quicker reactions in both lane-change and color discrimination tasks, especially in scenarios with longer time intervals between stimuli. This suggests that the order of tasks plays a role in how music affects performance in dual-task situations. Our findings also confirmed previous research showing that presenting a stimulus before a driving task increases RT. Furthermore, music positively influenced driving accuracy, potentially by inducing positive emotions and creating a controlled acoustic environment. While music tempo had little effect on lane-change performance, it significantly influenced cognitive processing speed in the color discrimination task, without impacting accuracy. Importantly, participants’ perceived mental workload remained consistent across all conditions. These insights shed light on how background music choice can enhance driving performance, potentially contributing to safer and more precise driving experiences.

## Data Availability

The datasets used and/or analyzed during the current study available from the corresponding author on reasonable request.

